# Patterns of annual and seasonal immune investment in a temporal reproductive opportunist

**DOI:** 10.1101/651158

**Authors:** Elizabeth M. Schultz, Christian E. Gunning, Jamie M. Cornelius, Dustin G. Reichard, Kirk C. Klasing, Thomas P. Hahn

## Abstract

Historically, investigations of how organisms’ investments in immunity fluctuate in response to environmental and physiological changes have focused on seasonally breeding organisms that confine reproduction to seasons with mild environmental conditions and abundant resources. Consequently, knowledge of how harsh environmental conditions and reproductive effort may interact to shape investment in immunity remains limited. The red crossbill, *Loxia curvirostra*, is a songbird that can breed on both short, cold and long, warm days if conifer seeds are abundant. This species provides an ideal system to investigate the influence of environmental fluctuations, reproductive investment, and their potential interactions on patterns of immune investment. In this study, we measured inter- and intra-annual immune variation in crossbills across four consecutive summers (2010-2013) and multiple seasons within one year (summer 2011-spring 2012) to explore how physiological and environmental factors impact this immune variation. Overall, the data suggest that immunity varies seasonally, among years, and in response to environmental fluctuations in food resources, precipitation, and temperature, but less in response to physiological measures such as reproduction. Collectively, this system demonstrates that a reproductively flexible organism may breed when conditions allow simultaneous investment in survival-related processes rather than at the expense of them.

## Introduction

Many temperate, terrestrial organisms experience extensive seasonal variation in weather, disease potential, and resource availability across the annual cycle. In turn, natural selection favours strategies that balance seasonal allocation to both reproduction and self-maintenance to maximize fitness [1]. Investment in immune function promotes survival by minimizing deleterious effects of pathogens and disease [2]. However, the energy and opportunity costs involved in maintaining immunity can be high [3, 4]. Empirical data suggest that changing environmental conditions (e.g., pathogens, resource availability) strongly influence allocation to immunity [5, 6]. If varying environmental conditions most strongly influence immune allocation, then investment in immunity would vary significantly both within and between years according to prevailing conditions (*sensu* Hegemann et al. [7]). However, organisms may also modulate immunity in direct response to an energy trade-off with competing processes such as reproduction, migration, or plumage/pelage moult. These trade-offs would result in consistent, cyclical patterns of immune investment that would vary little between years (*sensu* Hegemann et al. [7]).

Effects of varying environmental conditions and competing physiological processes on immune investment are not mutually exclusive, and our ability to quantify the relative contributions of these effects has been limited by both experimental methodology and study systems. Previous research on seasonal variation in immunity has focused on small mammals [8], with studies of birds focusing only on single stages of the annual cycle [6], and generally finding that investment in immunity decreases during reproduction (e.g., [9, 10]), moult [11, 12], migration [13–15], and winter [16, 17], but there are notable exceptions to this trend [6, 18]. A few studies have examined modulations in immunity across multiple annual cycle stages. Whereas components of immunity like microbial-killing ability were reduced during breeding in house sparrows (*Passer domesticus*) [19], complement activity, natural antibody levels, total immunoglobulin levels, and antibody response to multiple foreign antigens were higher during breeding in house sparrows [19], great tits (*Parus major*) [20], and skylarks (*Alauda arvensis*) [7], when compared to birds caught during moult and winter. These results suggest that significant seasonal variation in immune investment depends on concurrent investment in processes like reproduction and plumage moult. However, disentangling environmental from physiological effects on immune investment requires sampling across multiple years. The few studies that have examined interannual variation in immune investment demonstrated significant interannual variation in metrics of constitutive immunity (complement activity, natural antibody, and haptoglobin levels) in skylarks [7] and seven species of Galápagos finches [21], suggesting that annual variation in environmental conditions may contribute to variation in immunity. However, these studies of long-term temporal variation have not focused on causal mechanisms of the observed patterns.

Nearly all previous research on this topic has focused on seasonal breeders that perform most demanding annual cycle processes during high resource availability and benign environmental conditions, making it difficult to identify the relative importance of environmental and physiological factors in determining investment in immunity [5, 6]. As such, more studies are needed on free-living vertebrates across multiple seasons within a year, and across multiple years, to better understand the factors underlying seasonal differences in immunity. Further, investigating these questions in organisms whose reproduction is facultative across a wide range of environmental conditions will facilitate a more direct assessment of how physiological demands and environmental fluctuations influence the evolution of life history-related investments in immunity.

Here we present a multiannual study of a songbird, the red crossbill (*Loxia curvirostra*), that breeds both in summer and winter if food (i.e. conifer seeds) is sufficiently abundant [22]. We quantified the relative importance of environmental (e.g., ambient temperature and food availability), and physiological (e.g., breeding and integument moult) variables to immune investment and examined intra- and inter-annual variation in multiple measures of immunity in free-living red crossbills. Constitutive immunity was measured as an important first-line of defence against invading pathogens; its consistent production costs may underlie physiological trade-offs [6, 23]. Specifically, we quantified 1) complement and natural antibody levels via the hemolysis-hemagglutination assay [24]; 2) acute-phase protein concentrations (haptoglobin analog PIT54 in birds) [25]; and 3) the relative abundances of circulating leukocytes.

For model construction, we employed a two-tiered approach for each response. We first used a suite of random forest models (RFMs) for variable selection [26, 27], allowing us to identify putative predictors that warranted further exploration. The RFM is a non-parametric model formulation that embeds fewer assumptions than traditional linear models (LMs), facilitating estimation of nonlinear effects and high-order interactions. RFMs, however, lack the familiarity and robust inference framework of LMs. As such, we used the variables selected by each RFM to build a corresponding LM. We used the resulting LMs to examine the effect of these putative predictors, including environmental and physiological covariates, and other covariates known to affect immune variation in other systems (e.g., sex and age, reviewed in Adelman [18]). This approach allows us to highlight the relative contribution of temporal and environmental variables to annual and seasonal variation in immunity.

## Materials and Methods

### Field Methods

#### STUDY SPECIES AND SITE

Red crossbills (*Loxia curvirostra*) are nomadic, reproductively flexible passerines that eat mostly conifer seeds, the availability of which varies dramatically in space and time [22, 28, 29]. In years with abundant cone crops, crossbills can breed nine months of the year, with potentially multiple broods per year, despite thermal challenges and short days in some seasons [30–32]. Crossbills are categorized as “seasonal opportunists;” while they are more behaviourally flexible than seasonal migrants, they also exhibit seasonal cycles of migratory physiology, reproduction, and moult that are not principally predicted by variation in food supply [33–36].

Data were collected from free-living red crossbills from 2010-2013 around Grand Teton National Park, Wyoming, USA (N 43° 45’, W 110° 39’), a montane temperate-zone environment with large seasonal fluctuations in day length, food availability, temperature, and precipitation (Fig. 1A, C, D). Crossbills were sexed and aged using plumage and skull as described in Pyle [37]. At this site, red crossbill abundance fluctuates from year-to-year and is related to the cone crop size on dominant conifers [32, 38]. The dominant conifers used by crossbills here are lodgepole pine (*Pinus contorta*), Douglas-fir (*Pseudotsuga menziesii*), Engelmann spruce (*Picea engelmannii*), and blue spruce (*Picea pungens*). Ten described vocal “types” of red crossbills can be categorized into four classes by body size and bill morphology [39–41]. These morphological differences among types optimize their foraging efficiency on specific conifer taxa [39, 40, 42]. This study presents immunity data from vocal types 2, 3, 4, and 5 (Table S2).

**Figure 1:**
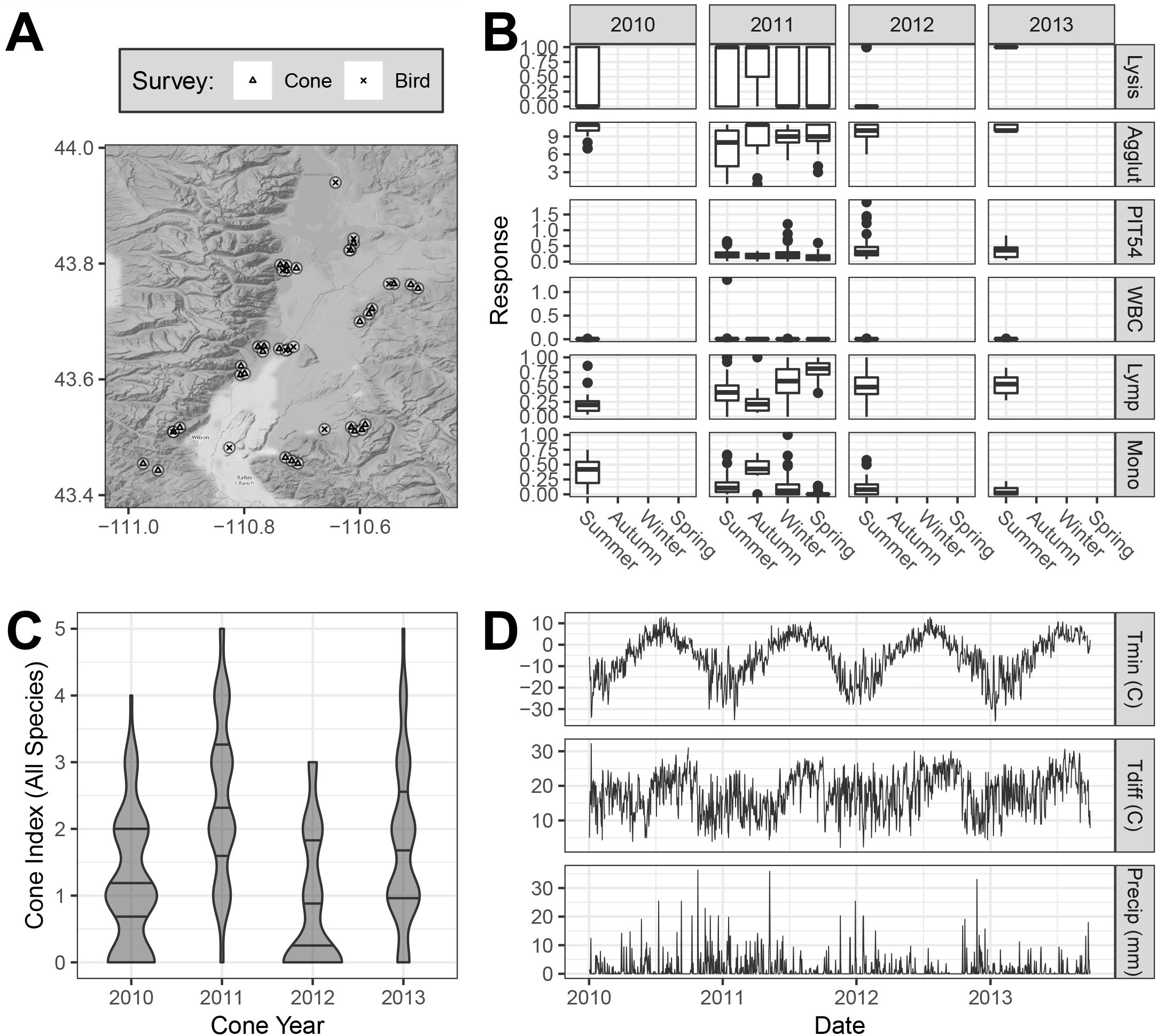
**A.** Map of Survey Sites (Teton County, Wyoming, USA). **B.** Distribution of each response over time. Column labels show cone year (June 1 of the current year through May 31 of the subsequent year). During each cone year, crossbills were sampled during (at least) the summer season; crossbills were also sampled in each season in cone year 2011 (June 1, 2011 - May 31, 2012). Responses: Lysis (non-zero hemolysis score), Agglut (agglutination score), PIT54 (mg/mL), WBC (Prop Leukocytes/Erythrocytes), Lymp (Prop Lymphocytes/Leukocytes), Mono (Prop Monocytes/Leukocytes). **C.** Violin plot of cone index by cone year (Score: 0-5) over all species (BS, DF, ES, and LPP; see Fig. S3 for details). Black horizontal lines show the Q1, median, and Q3. Width is proportional to sample count. **D.** Weather time series from Moose, Wyoming (GHCND Site ID: USC00486428). Precipitation shows liquid equivalent. For each weather covariate, a rolling mean was used for subsequent analysis (see Table S5).

#### DELINEATION OF SEASON AND CONE YEAR

Sampling periods were categorized into seasons: birds caught in the summer were caught from June 23 and September 12, autumn from October 25-30, winter from March 1-11, and spring from May 3-9. Sample sizes per year and season appear in Table S1. A “cone year” coincides with the cone development occurring between approximately June 1st of one year until the following spring when old cones are depleted or new cones start developing [43] (See below).

#### CAPTURE METHODS AND BLOOD SAMPLING

Crossbills were lured into mist nets with live caged decoys and/or playback. Approximately 300 μL of blood per bird was collected from the brachial vein into heparinized microhematocrit capillary tubes. This collection occurred between 0700 and 2000 hours with a median elapsed time from capture to sampling of 3.73 minutes (maximum of 60 minutes) to minimize potential effects of rising glucocorticoids [44–46]. Blood samples were held on ice for no more than seven hours before centrifuging (10 min at 10000 RPM, IEC clinical centrifuge) and separated plasma was stored at −20°C until immune assays were performed. Percent packed cell volume (hematocrit) was measured in all birds except those captured during summer 2010.

### Physiological Measures

#### REPRODUCTIVE MEASURES

Male cloacal protuberance (CP) length was measured with dial callipers from the abdomen to the cloacal tip; this measure correlates to testis length in free-living male red crossbills [33]. For females, the brood patch was scored as 0 (a dry, fully feathered breast), 1 (loss of feathers, no vascularization), 2 (loss of feathers with mild oedema and/or vascularization), 3 (loss of feathers, full oedema/vascularization), or 4 (bare and wrinkly breast, post full oedema); brood patch scores of >0 significantly predict crossbill ovary condition [33].

#### PLUMAGE MOULT INTENSITY

Pre-basic moult occurs seasonally in red crossbills (June-November) and may be arrested during summer breeding [22, 35]. The primary flight feathers grow sequentially from wrist to wingtip [47], and the number of actively growing feathers was defined as primary or flight feather moult intensity. Contour feather (body) moult was scored by surveying the entire body and rated on a scale of 0 (no pins or sheaths present), 1 (light: few pins, growing or sheathed feathers in one tract), 2 (medium: ~10-20 pins, growing or sheathed feathers in multiple tracts), and 3 (heavy: many pins, growing/sheathed feathers across multiple feather tracts) [47].

#### MASS, FAT, AND STRUCTURAL MEASURES

Body mass was measured to the nearest 0.1 grams with a Pesola spring scale and furcular and abdominal fat were scored on a scale from 0 (no fat) to 5 (bulging) [48, 49]. Tarsometatarsus was measured using dial callipers (in millimetres) and body condition was calculated by performing linear regressions of mass by tarsus length and calculating residuals.

### ENVIRONMENTAL MEASURES

#### CONE CROP

To evaluate the availability of conifer food sources in the area (lodgepole pine, Douglas-fir, Engelmann spruce, and blue spruce), one observer (TPH) visited 12 distinct, long-term point-count sites between July and September of each year (Fig. 1A). These sites were established in 2006, and at each annual visit, 10-20 trees of each species present within 50 meters of the point-count site were assigned a cone abundance index (USFS 1994) ranging from 0-5 (0 no cones) to 5 (abundant cones on cone-bearing section of tree) [38]. Seed supply is greatest in summer and early autumn when cones are ripening and beginning to open. Seed abundance declines in winter and spring due to seed shedding and predation [50].

#### LOCAL WEATHER CONDITIONS

For each day of bird capture, 24-hour precipitation amounts (mm) and daily maximum and minimum temperature (degrees Celsius) were accessed from the National Oceanic & Atmospheric Administration (NOAA) National Climate Data Centre website, using the weather station MOOSE 1 NNE, Wyoming, USA (Elevation: 1970.84 meters, Latitude: 43.662° N Longitude: 110.712° W) (Fig. 1D). By subtracting the daily minimum (Tmin) from maximum temperatures (Tmax), we calculated the temperature difference (Tdiff) for each capture day to compute diurnal temperature range, which can vary substantially in a montane environment. Minimum and maximum daily temperature were highly correlated; we thus omitted maximum daily temperature and only included Tmin, which in this climate would have a greater impact on thermoregulatory demand, and Tdiff for subsequent analyses.

As organisms can respond to environmental conditions over a range of time scales, we also computed rolling means for each weather variable over a range of ecologically plausible window lengths (R package *zoo,* rollapply function, right-aligned, NAs omitted). For each window length and combination of (predictor x weather), the Spearman correlation coefficient was computed (Table S5, Fig. S9). For each weather predictor and window length, (absolute) correlation coefficients were averaged across responses; the window length with the maximum mean correlation was used for subsequent analysis (Table S5). This method allowed us to account for delayed response between environmental drivers and ecological responses, while maintaining a parsimonious and non-parametric approach to data processing and variable selection.

### IMMUNE ASSAYS

#### COMPLEMENT AND NATURAL ANTIBODIES (Lysis and Agglutination)

The protocol described in Matson et al. [24] was used to measure plasma complement activity and non-specific natural antibodies via red blood cell lysis and agglutination, respectively. Partial lysis and agglutination were indicated by half scores. Samples were scored blind to sampling date and randomized across plates by one observer (EMS). A positive standard (chicken plasma) was run on all plates in duplicate. 10 μL of plasma was used (instead of 25 μL) due to small blood volumes and reagent volumes were adjusted accordingly [51]. The average inter-plate variation (standard deviation) was 0.28 lysis titres and 0.09 agglutination titres. Due to the abundance of lysis scores of zero in our dataset (60.7% zero scores; non-zero scores ranged from 0.4-5), we assigned individuals a 0 or 1 score, where 1 was any non-zero lysis score.

#### HAPTOGLOBIN (PIT54)

To quantify plasma PIT54 concentrations, a colorimetric assay kit (TP801; Tri-Delta Diagnostics, NJ, USA) was used. To accommodate small blood volumes, 5 μL was used (instead of 7.5 μL) and all reagents were adjusted accordingly[51]. Because additional plasma was used to optimize the hemolysis-hemagglutination assay in 2010, haptoglobin values were not measured.

#### CIRCULATING CELLULAR IMMUNITY (WBC)

To identify the quantity and type of leukocytes (lymphocytes, heterophils, monocytes, eosinophils, and basophils), a drop of whole blood was spread onto a slide, air-dried, fixed with 100% methanol, and stained with Wright-Giemsa (Cambridge Diagnostic Camco Stain Pack). The number and type of leukocytes under 1000x magnification were scored by two observers using the methodology outlined in Campbell [52]: EMS scored all seasons except for summer 2012 which was done by D. Jaul, who was trained by and calibrated against EMS. Leukocytes were detected across 100 microscope “fields” or approximately 10,000 erythrocytes and reported as the number of leukocytes/number of fields scored. We also calculated the heterophil to lymphocyte ratio [53], and the relative proportion of the each leukocyte type [54]. Eosinophil, heterophil, basophil, and heterophil to lymphocyte ratio models had poor predictive value; we only report results from overall leukocytes (WBC), lymphocytes (**Lymp**) and monocytes **(Mono).**

### Statistical Analyses

#### OVERVIEW

All analysis was conducted in R [55]. We constructed a set of statistical models for each of six separate immune measures (**Lysis, Agglut, PIT54**, **WBC, Lymp, Mono); immune responses, except for Mono and Lymp, were not highly correlated see Fig. S7**), using separate models for seasonal and annual variation (**Season, Year)**. Model predictors included the following variables (Table S3): environmental (cone year, rolling-mean of daily minimum temperature (Tmin), Tdiff, and precipitation (Precip)); physiological (CP length/BP score, primary and contour feather moult intensity, hematocrit score, residual body mass score, and composite fat score); intrinsic (age, sex, vocal type); and sampling-related (capture location, time of day, and time elapsed between capture and blood sampling). We first used random forest models for variable selection, and then constructed linear models for statistical inference using the previously selected variables [56].

#### STATISTICAL MODELS

Owing to data limitations, we separated data sets into two groups based on observation timing: yearly (summer, cone years 2010-2013) and seasonal (summer and fall 2011, winter and spring 2012 i.e. cone year 2011); see “Delineation of Season” in Methods for date ranges. We constructed models within each group, including sampling period (cone year or season, respectively) as a predictor in the relevant model.

For each combination of (time x measure), we constructed an unbiased random forest model (RFM) (R package party, [57–59]) including all measured covariates (Table S3). For each RFM, we computed variable importance (mean decrease in accuracy after permutation).We identified covariates with negative variable importance and removed them from further consideration, and then refit the model using the remaining covariates. We repeated this process three times, so that the remaining predictor variables show a consistent statistical association with the given immune measure. An adjusted goodness-of-fit (R^2^) of each final RFM was assessed by first predicting the model’s response using out-of-bag (OOB) observations; sum of squared residuals (SS_Resid_) were computed, with R^2^ = 1−(SS_Resid_/SS_Total_), which was then adjusted for sample size and number of covariates using an approximate Wherry formula (Table S4).

For each combination of time x immune measure, we then constructed a linear model (LM) that including all covariates with positive variable importance from the final RFM (above). A logistic GLM was used to model Lysis (0= no hemolysis, 1= non-zero hemolysis), which we refer to simply as the Lysis LM. For each LM, we report Type II ANOVAs for each predictor, and an overall model p-value.

Due to multicollinearity among variables in the model (Fig. S8), coefficient p-values and standard errors are generally considered biased and unreliable [61–63]. Nonetheless, overall model significance, R^2^ values, and coefficients are interpretable within the context of the model [64, 65]. As such, we intentionally avoid variable selection on the basis of p-values [66], and instead report for each immune measure and time period a full model including all covariates selected by the corresponding RFM.

To assess the direction and magnitude of response to temporal variation, we computed expected marginal means (EMM) by time period for each immune measure, both for yearly and seasonal LMs (R package “*emmeans*”[60]). For each LM, we also summarize model coefficient summaries for all continuous predictors with p <0.2.

## Results

Hematocrit, vocal type, and all sampling-related variables (capture location, time of day, and time elapsed between capture and blood sampling) had negative variable importance for all immune parameter RFMs and thus were eliminated from subsequent LMs due to their negligible predictive value. Overall variable importance varied widely between models; for yearly models, only cone year was consistently high, while for seasonal models variable importance was consistently positive only for Tmin and season. For both yearly and seasonal models moult, covariates with consistently low variable importance include fat, reproductive condition, sex, and age (Fig. S1).

Overall fits of final models (i.e., adjusted R^2^s) were generally higher for LMs than RFMs (note, however, that RFM results are not directly comparable to LMs), though their magnitudes broadly agree within each response and time (Table S4). WBC displayed consistently poor fits for both yearly and seasonal models; the yearly PIT54 model also displayed poor fit (Table S4). Yearly LMs displayed relatively high R^2^s for Lysis (0.32), Agglutination (0.27), and Monocytes (0.27); for seasonal LMs, Lymphocytes (0.2) and Monocytes (0.21) exhibit relatively high R^2^s (Table S4).

### Interannual variation

Cone year had the highest variable importance in all yearly RFM models (Fig. S1) and the corresponding LMs revealed cone year as a significant predictor of annual variation in complement (Lysis), natural antibodies (Agglut), PIT54, total leukocytes (WBC), lymphocytes (Lymp), and monocytes (Mono) (Tables S7-12).

In addition to cone year, other covariates with positive variable importance within annual RFMs include age, sex, reproductive condition (CP length and BP scores), temperature difference (Tdiff, 8 days prior to and including day of capture), minimum temperature (Tmin, within a day of capture), precipitation (Precip, 32 days prior to and including day of capture), body condition (R.mass), body and flight feather moult intensity, and fat scores (Fig. S1). In the LMs, flight feather moult (Ff) and Tdiff were identified as important predictors of annual variation in complement and total WBC, respectively (Table S6-7, S12).

Cone crop scores varied significantly among tree species across cone year (see Figs 1C, S3, and S4 for details). Summaries of immune measures and corresponding LM expected marginal means (EMM) indicated higher complement and total leukocyte counts in cone years with larger cone crops (2011, 2013) and lower levels in cone years with smaller crops (2010, 2012) (Figs. 1B, 2A, S5). In contrast, natural antibodies and PIT54 were negatively related to cone crop, with lower scores in 2011 and the highest scores in cone years with smaller crops (2012, 2013) (Figs. 1B, Fig. 2A, S5). The relationship between cone crop and monocytes and lymphocytes was not linear; highest and lowest levels of lymphocytes and monocytes, respectively, occurred in 2013, a mid-level cone year (Figs 1B, 2A, S5).

**Figure 2:**
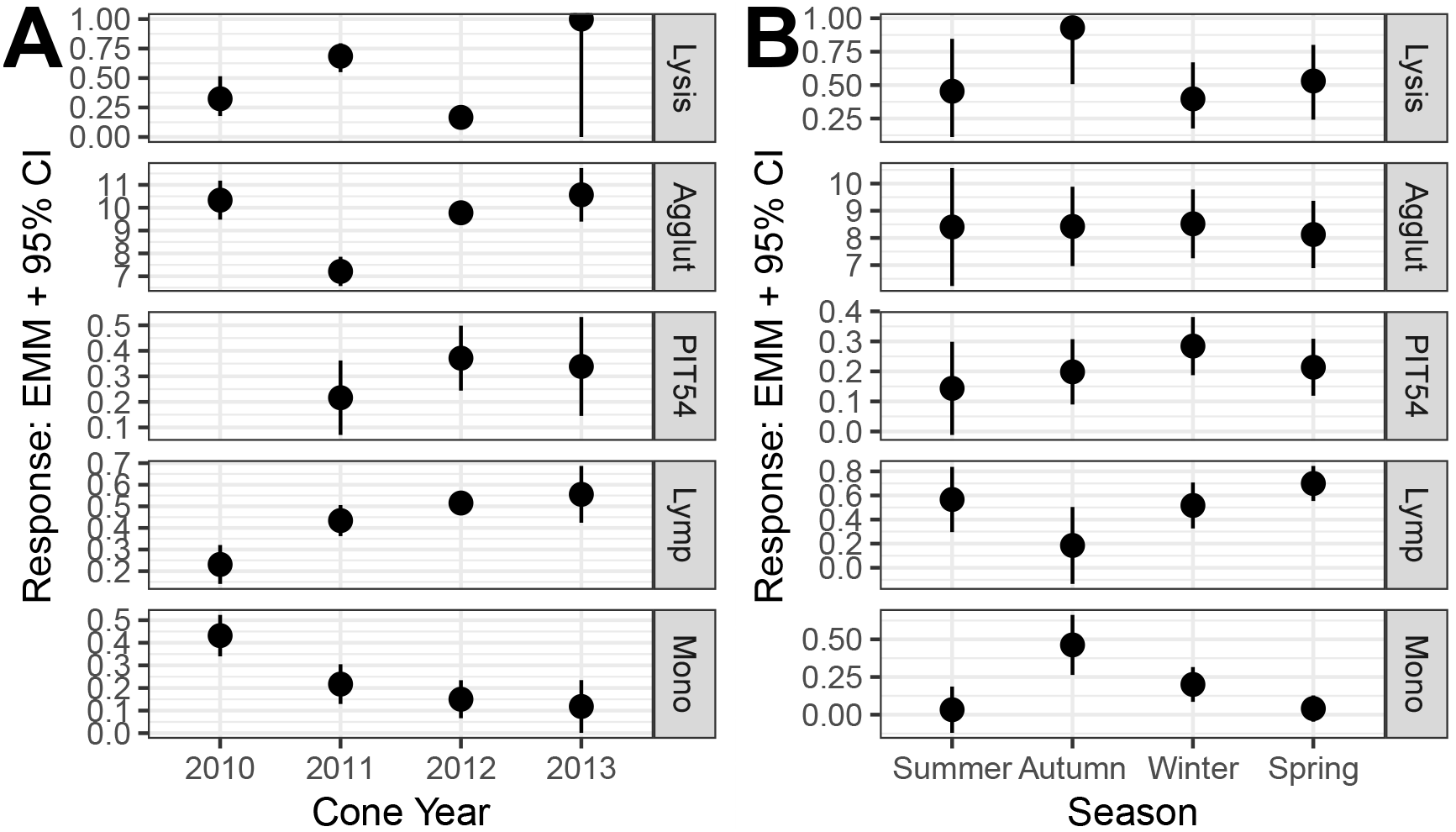
Expected marginal mean (EMM) and 95% CI of each response by time period: **A.** Cone Year (all years, Summer only). LM R^2^: Lysis=0.32, Agglut=0.27, PIT54=0.05, Lymp=0.17, Mono=0.27, WBC=0.01. **B.** Season (all seasons, Cone Year 2011 only). LM R^2^: Lysis=0.14, Agglut=0.09, PIT54=0.13, Lymp=0.20, Mono=0.21, WBC=0.02. Models with R^2^ < 0.05 are omitted for clarity (see also Table S4).

### Seasonal variation

Season consistently showed positive variable importance in RFMs predicting total leukocytes, lymphocytes, monocytes, natural antibodies, PIT54, and complement (Fig. S1), Monocytes and lymphocytes displayed the most substantial seasonal fluctuations, with extremes in autumn, when lymphocytes were the lowest and monocytes the highest (Fig. 2B, Tables S10-11).

Ambient temperature (Tdiff, Tmin), precipitation, body condition, contour and flight feather moult had positive variable importance in the RFM predicting seasonal variation in complement (Fig. S1). Of these variables, Tdiff, Precip, and R.mass were identified in the LM as being most related to complement (Tables S6-7). Specifically, these variables were positively related, suggesting that crossbills had higher complement when they were in better condition, or precipitation or ambient temperature were higher. Ambient temperature (Tmin and Tdiff), body fat, and reproductive condition had positive variable importance in the RFM predicting seasonal variation in natural antibodies (Fig. S1). In the corresponding LM, only Tdiff was weakly, negatively related to natural antibodies within the LM (Tables S6, S8). Precipitation, body condition, fat, and ambient temperature (Tmin, Tdiff) had positive variable importance in the RFM predicting seasonal variation in PIT54 (Fig. S1). Precipitation, body condition, and Tdiff were most related to PIT54 in the LM (Tables S6, S9). These variables were positively related, suggesting that birds in better body condition or sampled subsequent to period of greater precipitation, temperature have higher PIT54 concentration (Table S6).

Precipitation, ambient temperature (Tdiff, Tmin), flight feather and body moult, sex, age, reproductive condition, and body condition all had positive variable importance in RFMs predicting seasonal variation in total leukocytes, lymphocytes, and monocytes (Fig. S1). While only Tdiff was a substantial contributor to seasonal WBC variation (Table S6, S12), ambient temperature (Tdiff, Tmin), precipitation, and flight feather moult rank highly in LMs predicting seasonal variation in monocytes and lymphocytes (Table S6, S10, S11). Taken together, temperature was positively related to total leukocytes and monocytes, whereas precipitation and temperature were negatively related to lymphocytes.

## Discussion

We observed significant interannual variation in natural antibodies, complement, total leukocytes, monocytes, lymphocytes, and PIT54 concentrations among four consecutive summers (2010-2013) (Figs 1, 2A), however model fit for PIT54 and total leukocytes was poor (Table S4). Concurrently, the size of the cone crops of important conifer species (Figs 1C, S3-4), which are primary food resources for crossbills, demonstrated significant annual variation, that may be driven, in part, by past climatic conditions [67, 68]. This annual variation in cone crop during our study corresponded to the modulation of several immune measures. Complement and total leukocytes were higher when cone crops were larger, whereas natural antibodies and acute phase proteins (PIT54) were higher when cone crops were smaller. Higher levels of natural antibodies when cone crops are small may reflect investment in “low-cost” immunity when resources are limiting [69]. In contrast, higher levels of PIT54 and lower levels of other immune measures (i.e. complement and total leukocytes) when food resources are low could indicate a trade-off within the immune system to favour higher-cost, more rapid immune defence [70]. Further, this suggests that the overall costs of constitutively maintaining higher levels of protective proteins and cells is higher than occasionally inducing an expensive inflammatory response. While PIT54 was, on average, lower in cone years with large cone crops, it was higher in lower cone years, which could alternatively reflect an activated immune system due to infection or inflammation. However, because we do not have repeated samples from individuals, it is difficult to ascertain if the higher levels of PIT54 indicate inflammation or are actually within the normal range of the individual.

Seasonal variation in crossbill immune activity was detected by examining modulations within birds sampled across a single cone year (summer 2011-spring 2012), corresponding to the entire annual cycle of one large, cumulative cone crop. Within this year, there was seasonal variation in natural antibodies, complement, PIT54, and total leukocytes, monocyte, and lymphocytes (Figs 1B, 2B), however model fit for total leukocytes was poor (Table S4). Ambient temperature and precipitation were consistent predictors of this variation: larger differences in minimum and maximum daily temperature (Tdiff) were positively related to complement, PIT54, and total leukocytes and negatively related to natural antibodies. Alternatively, this overall increase in immune investment could be linked to a higher probability of disease and infection during the summer months, when Tdiff was overall higher. For example, red crossbills have higher *Haemoproteus* infections in the late spring and summer than other times of year [43, 71], and this was significantly related to total leukocyte counts [43].

Cumulative exposure to precipitation, when measured 32 days prior to and including day of capture, was strongly and positively related to PIT54. This may be driven, in part, by higher precipitation in the form of snowfall during winter months (Fig. S2). Higher levels of the acute phase C-reactive protein have been found in human populations during winter months [72], but it is difficult to determine the precise mechanism explaining this relationship between snowfall and PIT54. It is possible that higher PIT54 in crossbills indicated an activated immune system due to increased risk of infection or inflammation that was due to or exacerbated by a challenging thermal and foraging winter environment [73]. In support of an activated immune system, complement was also positively related to precipitation. However, in contrast, body condition related positively to PIT54, suggesting that either PIT54 is not reflective of immune challenge or infection did not cause reduced body condition. PIT54 was also higher during years of low cone supply and may be a favoured immune investment when resources are limited or conditions are challenging. Evidence pointing to a relationship between condition and immune investment in free-living organisms is mixed, with research suggesting no relationship between body condition and specific antibody response in mallards (*Anas platyrhnchos*)[74], and other work showing that elk (*Cervus elaphus*) in poor nutritional condition invest more in constitutive immune measures [75].

Although “costly” physiological processes such as reproduction and moult can affect immune investment [9, 11, 76–78], our data only weakly supported this prediction. While both plumage moult intensity and reproductive measures were included in RFMs and LMs predicting complement, total leukocytes, monocytes, lymphocytes, and natural antibodies, only flight feather moult displayed any strong relationship with interannual or seasonal variation. Specifically, flight feather moult was positively related to annual variation in complement, and seasonal variation in monocytes. In temperate breeding, northern hemisphere birds, flight and contour moult are generally heaviest during autumn (Fig. S6,[47]), which in crossbills corresponded to the lowest and highest lymphocyte and monocyte proportions, respectively. Elevated monocyte concentration during moult has also been documented in red knots (*Calidris canutus*) [79]. Given that feathers erupting through the skin can induce dermal inflammation [80], and monocytes are phagocytic leukocytes integral to the inflammatory response [52], elevated monocytes during peak moult is not surprising. The decrease in lymphocytes during this same period may indicate a trade-off within the immune system to favour phagocyte-mediated responses rather than cellular and antibody-based immunity. Overall, we are somewhat limited by our sample size during this period: capturing crossbills during moult is especially difficult because more secretive behaviours accompany this stage [81], despite our large field effort.

Although our models did not identify reproduction as a good predictor of annual or seasonal immune variation, we note that the reproductive covariates measured and described here do not represent the total investment or cost of reproduction. Although CP length and BP score are correlated with testes length and ovary condition in crossbills [33], these measures do not quantify the total reproductive costs incurred by females and males (e.g., egg laying and incubation, provisioning offspring), which are energetically demanding in birds [82]. Reproduction should be even more demanding for crossbills rearing young in winter due to higher energy and thermoregulatory demands. Breeding crossbills in winter show intensive foraging behaviour to meet the combined energy demands of themselves and their incubating mates and growing nestlings (J.M.C. unpublished radio-tracking data). Baseline corticosterone levels, which reflect current energy demands [83], are not higher in winter-breeding crossbills, nor are they higher in breeding crossbills in general [33]; however male heart-rate telemetry data during this period demonstrate elevated metabolic rate (J.M.C. unpublished data). Despite elevated demands, crossbills likely cope with the high cost of winter reproduction in part because reproduction occurs in winter only if conifer seeds are abundant and foraging efficiency is high, which may allow for investment in immunity. However, at least some crossbills still attempt reproduction in low cone years (Fig. S6). Finally, crossbills may incur lower overall reproductive costs compared with other seasonal species because they do not defend territories and have relatively small clutch sizes [22].

While our study highlights several proximate causal factors for immune investment, it is likely that some of this variation is in response to changes in photoperiod: a previous experimental study in captive red crossbills found that exposure to long days is sufficient to increase total leukocyte counts and bacterial killing ability [51]. In addition, our characterization of inter- and intra-annual variation in immunity is based on sampling from exclusively summer months and within one cone year, respectively. This sampling limitation means we are unable to fully tease apart environmental and physiological contributors to immunity, particularly because physiological covariates like reproduction, moult, and condition may vary significantly between and within years. In our study, reproduction, fat, flight feather moult, body moult, and condition did vary between and within years, although not substantially (Fig. S6). Finally, although we did not detect any significant immune variation among crossbill vocal types, our samples were comprised of multiple vocal types and thus potentially different populations of crossbills that could have experienced different environments and pathogens either prior to or after their arrival at our study site.

Overall, this study supports previous findings that 1) birds seasonally modulate investment in immunity [19, 20] and 2) investment in immunity changes between years, which has only been documented in two other studies—one on skylarks [7] and one on Galápagos finches [21]. Additionally, this study suggests that investment in immune function (i.e. complement-mediated lysis, antibody-mediated agglutination, haptoglobin/PIT54 concentrations, total leukocytes, monocytes, and lymphocytes) is related to changes in food resources, ambient temperature, precipitation, and plumage moult, and appears to be less related to investment in reproduction or other factors such as age or sex. This supports the hypothesis that crossbills breed when conditions allow simultaneous investment in survival-related processes rather than at the expense of them and agrees with behavioural ecology theories concerning the use of rich patches by highly mobile birds such as crossbills [84]. Finally, this study highlights the need for future studies to sample across multiple years and seasons in order to draw robust conclusions about the seasonality of immunity in wild organisms.

## Ethics

All bird capture and handling protocols were approved by the University of California Davis Institutional Animal Care and Use Committee (protocol number: 16729), US Federal Bird Banding Permit (22712), Wyoming Game & Fish Department (393), and Grand Teton National Park (GRTE-2010-SCI-0004,GRTE-2011-0005SCI, GRTE-2012-SCI-0004, GRTE-2013-SCI-0006).

## Data and Code Accessibility

For review, data files and R code available at Data Dryad (https://datadryad.org/review?doi=doi:10.5061/dryad.0fh3jn0) [85].

## Authors’ Contributions

EMS performed the immune assays, fieldwork, and wrote the manuscript; CEG and EMS carried out the statistical analyses, prepared the figures and tables, and edited the manuscript; JMC and DGR participated in fieldwork and contributed to the editing of the manuscript; KCK provided resources and laboratory space for immune assays and participated in editing of the manuscript; TPH provided partial funding, contributed fieldwork, and participated in editing of the manuscript. All authors approved the manuscript.

## Acknowledgements

We thank SE Knox, DZ Jaul, RE Koch, C Lopez, M Lohuis, and V Iseri for their help with data collection. We would also like to thank the UW-NPS Research Station and the Murie Center for providing housing and support during field collection.

## References

1. King J.R. 1974 Seasonal allocation of time and energy resources in birds. Cambridge, MA, Nuttall Ornithological Club.

2. Janeway C.A., Travers P., Walport M., Shlomchik M. 2004 Immunobiology: The Immune System in Health and Disease. Garland, New York.

3. Hasselquist D., Wasson M.F., Winkler D.W. 2001 Humoral immunocompetence correlates with date of egg-laying and reflects workload in female tree swallows. Behavioral Ecology 12(93-97).

4. Martin L.B., II, Scheuerlein A., Wikelski M. 2003 Immune activity elevates energy expenditure of house sparrows: a link between direct and indirect costs? Proceedings of the Royal Society B 270, 153–158.

5. Nelson R.J., Demas G.E. 1996 Seasonal changes in immune function. Q Rev Biol 71(4), 511–548.

6. Martin L.B., Weil Z.M., Nelson R.J. 2008 Seasonal changes in vertebrate immune activity: mediation by physiological trade-offs. Philosophical Transactions of the Royal Society B: Biological Sciences 363(1490), 321.

7. Hegemann A., Matson K.D., Both C., Tieleman B.I. 2012 Immune function in a free-living bird varies over the annual cycle, but seasonal patterns differ between years. Oecologia 170(3), 605–618.

8. Nelson R.J., Demas G.E., Klein S.L., Kriegsfeld L.J. 2002 Seasonal patterns of stress, immune function and disease. Cambridge, Cambridge University Press.

9. Ardia D.R. 2005 Tree swallows trade off immune function and reproductive effort differently across their range. Ecology 86(8), 2040–2046.

10. Bonneaud C., Mazuc J., Gonzalez G., Haussy C., Chastel O., Faivre B., Sorci G. 2003 Assessing the Cost of Mounting an Immune Response. The American naturalist 161(3), 367–379.

11. Ellis V.A., Merrill L., Wingfield J.C., O’Loghlen A.L., Rothstein S.I. 2012 Changes in immunocompetence and other physiological measures during molt in Brown-headed Cowbirds (*Molothrus ater*). The Auk 129(2), 231–238.

12. Martin L.B. 2005 Trade-offs between molt and immune activity in two populations of house sparrows (*Passer domesticus*). Canadian Journal of Zoology 83, 780–787.

13. Nebel S., Bauchinger U., Buehler D.M., Langlois L.A., Boyles M., Gerson A.R., Price E.R., McWilliams S.R., Guglielmo C.G. 2011 Constitutive immune function in European starlings, *Sturnus vulgaris*, is decreased immediately after an endurance flight in a wind tunnel. Journal of Experimental Biology 215(2), 272–278.

14. Owen J.C., Moore F.R. 2008 Swainson’s thrushes in migratory disposition exhibit reduced immune function. Journal of Ethology 26, 383–388.

15. van Gils J.A., Munster V.J., Radersma R., Liefhebber D., Fouchier R.A., Klaassen M. 2007 Hampered foraging and migratory performance in swans infected with low-pathogenic avian influencza A virus. PLoS ONE 2, 184.

16. Owen-Ashley N.T., Wingfield J.C. 2006 Seasonal modulation of sickness behavior in free-living northwestern song sparrows (*Melospiza melodia morphna*). Journal of Experimental Biology 209(16), 3062–3070.

17. Svensson E., Råberg L., Koch C., Hasselquist D. 1998 Energetic stress, immunosuppression and the costs of an antibody response. Functional Ecology 12(6), 912–919.

18. Adelman J.S., Ardia D.R., Schat K.A. 2013 Ecoimmunology. In Avian Immunology (eds. K.A. Schat, B. Kaspers, Kaiser P.), pp. 391–411, 2nd ed. Amsterdam, Elsevier Ltd.

19. Pap P.L., Czirják G.Á., Vágási C.I., Barta Z., Hasselquist D. 2010 Sexual dimorphism in immune function changes during the annual cycle in house sparrows. Die Naturwissenschaften 97(10), 891–901.

20. Pap P.L., Vágási C.I., Tökölyi J., Czirják G.Á., Barta Z. 2010 Variation in Haematological Indices and Immune Function During the Annual Cycle in the Great Tit Parus major. Ardea 98(1), 105–112.

21. Zylberberg M., Lee K.A., Klasing K.C., Wikelski M. 2012 Increasing avian pox prevalence varies by species, and with immune function, in Galápagos finches. Biological Conservation 153, 72–79.

22. Adkisson C. 1996 Red Crossbill. The Birds of North America 256, 1–23.

23. Schmid-Hempel P., Ebert D. 2003 On the evolutionary ecology of specific immune defence. Trends in Ecology & Evolution 18(1), 27–32.

24. Matson K.D., Ricklefs R.E., Klasing K.C. 2005 A hemolysis-hemagglutination assay for characterizing constitutive innate humoral immunity in wild and domestic birds. Developmental & Comparative Immunology 29(3), 275–286.

25. Matson K.D., Horrocks N.P.C., Versteegh M.A., Tieleman B.I. 2012 Baseline haptoglobin concentrations are repeatable and predictive of certain aspects of a subsequent experimentally-induced inflammatory response. Comp Biochem Phys A 162(1), 7–15.

26. De’ath G., Fabricus K. 2000 Classification and regression trees: a powerful yet simple technique for ecological data analysis. Ecology 81, 3178–3192.

27. Liaw A., Wiener M. 2002 Classification and Regression by randomForest. R News 2, 18–22.

28. Fowells H.A. 1968 Silvics of the United States. Washington, USDA Forest Service.

29. Koenig W.D., Knops J.M.H. 2000 Patterns of annual seed production by northern hemisphere trees: A global perspective. American Naturalist 155(59–69).

30. Benkman C.W. 1987 Food profitability and the foraging ecology of crossbills. Ecological Monographs 57, 251–267.

31. Benkman C.W. 1990 Intake rates and the timing of crossbill reproduction The Auk 107(2), 376–386.

32. Hahn T.P. 1998 Reproductive seasonality in an opportunistic breeder, the red crossbill, Loxia curvirostra. Ecology 79(7), 2365–2375.

33. Cornelius J.M., Breuner C.W., Hahn T.P. 2012 Coping with the extremes: stress physiology varies between winter and summer in breeding opportunists. Biology letters 8, 312–315.

34. Cornelius J.M., Hahn T.P. 2012 Seasonal pre-migratory fattening and increased activity in a nomadic and irruptive migrant, the Red Crossbill *Loxia curvirostra*. Ibis 154(4), 693–702.

35. Hahn T.P. 1995 Integration of photoperiodic and food cues to time changes in reproductive physiology by an opportunistic breeder, the red crossbill, *Loxia curvirostra* (Aves: Carduelinae). Journal of Experimental Zoology 272(3), 213–226.

36. Hahn T.P., Cornelius J.M., Sewall K.B., Kelsey T.R., Hau M., Perfito N. 2008 Environmental regulation of annual schedules in opportunistically-breeding songbirds: Adaptive specializations or variations on a theme of white-crowned sparrow? General and comparative endocrinology 157(3), 217–226.

37. Pyle P. 1997 Identification Guide to North American Birds. Part I. Columbidae to Ploceidae. Bolinas, CA, USA, Slate Creek Press.

38. Kelsey T.R. 2008 Foraging ecology, biogeography, and population dynamics of red crossbills in North America Davis, CA, University of California, Davis.

39. Benkman C.W. 2003 Divergent selection drives the adaptive radiation of crossbills. Evolution; international journal of organic evolution 57, 1176–1181.

40. Groth J.G. 1993 Evolutionary differentiation in morphology, vocalizations, and allozymes among nomadic sibling species in the North American red crossbill (Loxia curvirostra) complex. Berkeley, CA, University of California Press.

41. Irwin K. 2010 A new and cryptic call type of the red crossbill. Western Birds 41, 10–25.

42. Benkman C. 1993 Adaptation to single resources and the evolution of crossbill (Loxia) diversity. Ecological Monographs 63, 305–325.

43. Schultz E., Cornelius J., Reichard D., Klasing K., Hahn T. 2018 Innate immunity and environmental correlates of Haemoproteus prevalence and intensity in an opportunistic breeder. Parasitology, 1–12.

44. Buehler D.M., Bhola N., Barjaktarov D., Goymann W., Schwabl I., Tieleman B.I., Piersma T. 2008 Constitutive immune function responds more slowly to handling stress than corticosterone in a shorebird. Physiological and Biochemical Zoology 81(5), 673–681.

45. Matson K.D., Tieleman B.I., Klasing K.C. 2006 Capture Stress and the Bactericidal Competence of Blood and Plasma in Five Species of Tropical Birds. Physiological and biochemical zoology: PBZ 79(3), 556–564.

46. Millet S., Bennett J., Lee K.A., Hau M., Klasing K.C. 2007 Quantifying and comparing constitutive immunity across avian species. Developmental & Comparative Immunology 31(2), 188–201.

47. Cornelius J.M., Perfito N., Zann R.A., Breuner C.W., Hahn T.P. 2011 Physiological trade-offs in self-maintenance: Plumage molt and stress physiology in birds. Journal of Experimental Biology 214, 2768–2777.

48. Helms C.W.D., W. H. J.. 1960 Winter migratory weight and fat field studies on some North American buntings. Bird Banding 31, 1–40.

49. Nolan Jr V., Ketterson E.D. 1983 An analysis of body mass, wing length, and visible fat deposits of dark-eyed juncos wintering at different latitudes. Wilson J Ornithol 95(4), 603–620.

50. Burns R., Honkala B. 1990 Silvics of North America. In Agriculture Handbook (Washington, DC, U.S. Forest Service, Department of Agriculture.

51. Schultz E.M., Hahn T.P., Klasing K.C. 2017 Photoperiod but not food restriction modulates innate immunity in an opportunistic breeder, *Loxia curvirostra*. Journal of Experimental Biology 220, 722–730.

52. Campbell T.W. 1995 Avian Hematology and Cytology. Ames, IA, Iowa State University Press.

53. Davis A.K., Maney D.L., Maerz J.C. 2008 The use of leukocyte profiles to measure stress in vertebrate: a review for ecologists. Functional Ecology 22, 760–772.

54. Matson K.D., Cohen A.A., Klasing K.C., Ricklefs R.E., Scheuerlein A. 2006 No simple answers for ecological immunology: relationships among immune indices at the individual level break down at the species level in waterfowl. Proceedings Biological sciences / The Royal Society 273(1588), 815–822. (doi:10.1098/rspb.2005.3376).

55. R Core Team. 2018 R: A language and environment for statistical computing. (3.4.4 ed. Vienna, Austria, R Foundation for Statistical Computing.

56. Genuer R., Poggi J., Tuleau-Malot C. 2010 Variable selection using Random Forests. Pattern Recognition Letters 31(14), 2225–22236.

57. Hothorn T., Buehlmann P., Dudoit S., Molinaro A., Van Der Laan M. 2006 Survival Ensembles. Biostatistics 7(3), 355–373.

58. Strobl C., Boulesteix A., Kneib T., Augustin T., Zeileis A. 2008 Conditional Variable Importance for Random Forests. BMC Bioinformatics 9, 307.

59. Strobl C., Boulesteix A., Zeileis A., Hothorn T. 2007 Bias in Random Forest Variable Importance Measures: Illutrations, Sources and a Solution. BMC Bioinformatics 8, 25.

60. Lenth R. emmeans: Estimated Marginal Means, aka Least-Squares Means. (1.2.3 ed, R package

61. Vatcheva K., Lee M., McCormick J., Rahbar M. 2016 Multicollinearity in Regression Analyses Conducted in Epidemiologic Studies. Epidemiology (Sunnyvale) 6(2), 227.

62. Mason G. 1987 Coping with multicollinearity. The Canadian Journal of Program Evaluation 2, 87–93.

63. Mela C., Kopalle P. 2002 The impact of collinearity on analysis: the asymmetric effect of negative and positive correlations. Applied Economics 34, 667–677.

64. Smith A., Koper N., Francis M., Fahrig L. 2009 Confronting collinearity: comparing methods for disentangling the effects of habitat loss and fragmentation. Landscape Ecology 24, 1271–1285.

65. Mason C., Wd P.J. 1991 Collinearity, power, and interpretation of multiple regression analysis. Journal of Marketing Research 28, 268–280.

66. Amrhein V., Trafimow D., Greenland S. 2019 Inferential Statistics as Descriptive Statistics: There is No Replication Crisis if We Don’t Expect Replication. The American Statistician 73, 262–270.

67. Pearse S., Koenig D., Knops M. 2014 Cues versus proximate drivers: testing the mechanism behind masting behaviour. Oikos 123, 179–184.

68. Bisi F., von Hardenberg J., Bertolino S., Wauters L.A., Imperio S., Preatoni D.G., Provenzale A., Mazzamuto M.V., Martinoli A. 2016 Current and future conifer seed production in the Alps: testing weather factors as cues behind masting. European Journal of Forest Research 135(4), 743–754. (doi:10.1007/s10342-016-0969-4).

69. Lee K.A. 2006 Linking immune defenses and life history at the levels of the individual and the species. Integrative and comparative biology 46(6), 1000.

70. Buehler D.M., Encinas Viso F., Petit M., Vézina F., Tieleman B.I., Piersma T. 2009 Limited Access to Food and Physiological Trade-Offs in a Long-Distance Migrant Shorebird. II. Constitutive Immune Function and the Acute-Phase Response. Physiological and biochemical zoology: PBZ 82(5), 561–571.

71. Cornelius J.M., Zylberberg M., Breuner C.W., Gleiss A.C., Hahn T.P. 2014 Assessing the role of reproduction and stress in the spring emergence of haematozoan parasites in birds. Journal of Experimental Biology 217(6), 841–849.

72. Sung K. 2006 Seasonal variation of C-reactive protein in apparently healthy Koreans. International Journal of Cardiology 107, 338–342.

73. Owen-Ashley N.T., Wingfield, J.C. 2007 Acute phase responses of passerine birds: characterization and seasonal variation. J Ornithol 148 (suppl), S583–S591.

74. Arsnoe D., Ip H., Owen J. 2011 Influence of Body Condition on Influenza A Virus Infection in Mallard DUcks: Experimental Infection Data. PLoS ONE 6(8), e22633.

75. Downs C., Stewart K., Dick B. 2015 Investment in Constiutive Immune Function by North American Elk Experimentaly Maintained at Two Different Population Densities. PLoS ONE 10(5), e0125586.

76. Ben-Hamo M., Downs C., Burns D., Pinshow B. 2017 House sparrows offset the physiological trade-off between immune response and feather growth by adjusting foraging behavior. Journal of Avian Biology 48, 837–845.

77. Moreno J., Sanz J., Merino S., Arriero E. 2001 Daily energy expenditure and cell-mediated immunity in pied flycatchers while feeing nestlings: interaction with moult. Oecologia 129, 492–497.

78. Pap P., Vagasi C., Czirják G., Titilincu A., Pintea A., Barta Z. 2009 Carotenoids modulate the effect of coccidian infection on the condition and immune response in moulting house sparrows. Journal of Experimental Biology 212, 3228–3235.

79. Buehler D.M., Piersma T., Matson K., Tieleman B.I. 2008 Seasonal redistribution of immune function in a migrant shorebird: annual-cycle effects override adjustments to thermal regime. The American naturalist 172(6), 783–796. (doi:10.1086/592865).

80. Silverin B., Fange P., Viebke A., Westin J. 1999 Seasonal changes in mass and histology of the spleen in willow tits *Parus montanus*. Journal of Avian Biology 30, 255–262.

81. Morton G.A., Morton G.L. 1990 Dynamics of postnuptial molt in free-living mountain white-crowned sparrows. Condor 92, 813–828.

82. Monaghan P., Nager R.G. 1997 Why don’t birds lay more eggs? Trends in Ecology & Evolution 12, 270–274.

83. Romero L.M. 2002 Seasonal changes in plasma glucocorticoid concentrations in free living vertebrates. General and comparative endocrinology 128, 1–24.

84. Cornelius J.M., Watts H.E., Dingle H., Hahn T.P. 2013 Obligate versus rich patch opportunism: Evolution and endocrine mechanisms. General and comparative endocrinology 190, 76–80.

85. Schultz EM, Gunning CE, Cornelius JM, Klasing KC, Hahn TP. 2019. Data from: Patterns of annual and seasonal immune investment in a temporal reproductive opportunist. Dryad Digital Repository.(https://datadryad.org/review?doi=doi:10.5061/dryad.0fh3jn0)

