## Supplemental Tables and Figures for "Patterns of annual and seasonal immune investment in a temporal reproductive opportunist"

Table S1: Observations per time interval (cone year and season) for each response. Cone year columns (2010-2013) include only observations from summer. Season columns (Summer-Spring) include only observations from Cone Year 2011. Responses: Lysis (non-zero hemolysis score), Agglut (agglutination score), PIT54 (mg/mL), WBC (Prop Leukocytes/Erythrocytes), Lymp (Prop Lymphocytes/Leukocytes), Mono (Prop Monocytes/Leukocytes).

| Response | 2010 | 2011 | 2012 | 2013 | Summer | Autumn | Winter | Spring |
|---|---|---|---|---|---|---|---|---|
| Lysis | 29 | 55 | 96 | 13 | 55 | 11 | 98 | 26 |
| Agglut | 29 | 55 | 96 | 13 | 55 | 11 | 98 | 26 |
| PIT54 | 0 | 54 | 92 | 13 | 54 | 10 | 90 | 25 |
| WBC | 29 | 53 | 84 | 12 | 53 | 10 | 79 | 28 |
| Lymp | 29 | 53 | 84 | 12 | 53 | 10 | 77 | 27 |
| Mono | 29 | 53 | 84 | 12 | 53 | 10 | 77 | 27 |

Table S2: Summary of observations per time interval for each vocal type. The majority of observations were of vocal type 5 (72%), followed by type 2 (10.3%), type 4 (8.8%), unknown (NA, 4.7%), and type 3 (4.1%).

| Cone Year | Season | Type 2 | Type 3 | Type 4 | Type 5 | Type NA |
|---|---|---|---|---|---|---|
| 2010 | Summer | | | | 30 | |
| 2011 | Summer | 2 | 1 | 12 | 39 | 2 |
| 2011 | Autumn | | | | 11 | |
| 2011 | Winter | 4 | 2 | 11 | 74 | 8 |
| 2011 | Spring | 2 | 7 | | 19 | |
| 2012 | Summer | 23 | 4 | 6 | 63 | 5 |
| 2013 | Summer | 4 | | 1 | 8 | 1 |

Table S3: List of covariates selected by random forest models (RFM, see Figure S1). Weather variables were pre-processed to select one rolling mean window per covariate: Tmin = 1 day, Tdiff = 8 day, Precip = 32 day (see Table S5 for details). Excluded variables include site name, band id, capture time, capture duration, vocal type, and hematocrit. In addition, Tmax was excluded due to strong positive correlation with Tmin.

| Name | Units.and.Values | Notes |
|---|---|---|
| Cone.year | Integer: 2010-2013 | Cone growing season (start of year): June 1 through subsequent May 30 |
| Season | Category: Su, F, W, Sp | Astronomical season |
| Precip | mm | Total daily precipitation at Moose, WY |
| Tmin, Tmax | degrees C | Minimum and maximum daily temperature at Moose, WY |
| Tdiff | degrees C | Tmax - Tmin |
| Sex | Category: M, F | Determined from plumage |
| Age | Category: HY, AHY | Determined from plumage, skull: hatch year (HY) or after hatch year (AHY) |
| Body.molt | Integer: 0-3 | Score: contour feather moult intensity |
| CP/BP | Z-score (within sex) | Reproductive condition: cloacal protuberance length (M) or brood patch score (F) |
| Fat | Integer: 0-5 | Score: furcular and abdominal subcutaneous fat |
| Ff | Integer: 0-7 | Number of actively growing flight feathers (out of 18 total) |
| R.mass | grams | Body condition: tarsus mass regression residuals |

Table S4: Goodness-of-fit (adjusted $R^2$) for each model. Linear model (LM) adjusted $R^2$ were calculated using the `rsq.v` function in the R package `rsq` . Random Forest model (RFM): $R^2 = 1 - (SS_{resid}/SS_{total})$. Here, $SS_{total}$ was computed from the mean-only linear model, and residuals equal the difference between observed responses and out-of-bag RFM predictions. The RFM $R^2$ was adjusted using an (approximate) Wherry formula: adjusted $R^2 = 1 - (1 - R^2)(\frac{N-1}{N-V-1})$, for N=number of observations, V=number of covariates. RFM parameters: mtry=2; ntree=1000.

| Response | Year.RF | Year.LM | Season.RF | Season.LM |
|---|---|---|---|---|
| Lysis | 0.284 | 0.320 | 0.092 | 0.136 |
| Agglut | 0.275 | 0.267 | 0.062 | 0.095 |
| PIT54 | 0.024 | 0.054 | 0.058 | 0.131 |
| WBC | -0.057 | 0.008 | -0.075 | 0.015 |
| Lymp | 0.125 | 0.170 | 0.099 | 0.200 |
| Mono | 0.207 | 0.267 | 0.115 | 0.211 |

Table S5: Spearman correlation between each response and rolling mean of weather covariate over a range of window lengths (1 to 32 days, prior to and including crossbill sample date, NAs ommitted). Final windows were chosen to maximize mean absolute Spearman correlation over all responses: Tmin = 1 day ($|\bar{\rho}|$=0.19), Tdiff = 8 days ($|\bar{\rho}|$=0.18), Precip = 32 days ($|\bar{\rho}|$=0.12). See also Fig. S9.

| Response | Window | Tmin | Tdiff | Precip |
|---|---|---|---|---|
| Lysis | 1 | -0.079 | 0.040 | -0.101 |
| Lysis | 2 | -0.084 | 0.016 | -0.078 |
| Lysis | 4 | -0.112 | 0.045 | -0.107 |
| Lysis | 8 | -0.109 | 0.070 | -0.029 |
| Lysis | 16 | -0.088 | 0.062 | -0.011 |
| Lysis | 32 | 0.019 | 0.074 | 0.110 |
| Agglut | 1 | 0.177 | 0.025 | -0.032 |
| Agglut | 2 | 0.157 | 0.108 | 0.002 |
| Agglut | 4 | 0.162 | 0.129 | -0.012 |
| Agglut | 8 | 0.187 | 0.118 | 0.003 |
| Agglut | 16 | 0.206 | 0.136 | -0.059 |
| Agglut | 32 | 0.164 | 0.138 | -0.121 |
| PIT54 | 1 | 0.274 | 0.019 | 0.093 |
| PIT54 | 2 | 0.294 | 0.143 | 0.087 |
| PIT54 | 4 | 0.251 | 0.244 | -0.024 |
| PIT54 | 8 | 0.233 | 0.266 | -0.094 |
| PIT54 | 16 | 0.219 | 0.265 | -0.201 |
| PIT54 | 32 | 0.200 | 0.230 | -0.295 |
| WBC | 1 | 0.280 | 0.122 | 0.048 |
| WBC | 2 | 0.279 | 0.131 | 0.055 |
| WBC | 4 | 0.267 | 0.183 | 0.033 |
| WBC | 8 | 0.219 | 0.261 | -0.125 |
| WBC | 16 | 0.213 | 0.274 | -0.188 |
| WBC | 32 | 0.214 | 0.267 | -0.198 |
| Lymp | 1 | -0.177 | -0.044 | 0.000 |
| Lymp | 2 | -0.141 | -0.146 | 0.012 |
| Lymp | 4 | -0.131 | -0.177 | 0.108 |
| Lymp | 8 | -0.143 | -0.206 | 0.130 |
| Lymp | 16 | -0.153 | -0.125 | 0.109 |
| Lymp | 32 | -0.178 | -0.101 | -0.006 |
| Mono | 1 | 0.141 | -0.013 | 0.028 |
| Mono | 2 | 0.102 | 0.095 | -0.017 |
| Mono | 4 | 0.087 | 0.123 | -0.084 |
| Mono | 8 | 0.093 | 0.141 | -0.123 |
| Mono | 16 | 0.088 | 0.091 | -0.095 |
| Mono | 32 | 0.116 | 0.039 | 0.018 |

Table S6: LM regression coefficients. Horizontal lines show separate models (one model per time and response). Only continuous covariate coefficients (i.e., slopes) with p.value < 0.2 are shown. Within each model, estimates are ordered by p.value. Lysis: logistic GLM estimates back-transformed to linear scale, showing proportional change.

| Response | Time | Covariate | Est | SE | P.value |
|---|---|---|---|---|---|
| Lysis | Year | Ff | 1.28 | 0.137 | 0.071 |
| Lysis | Season | Tdiff | 1.24 | 0.0945 | 0.022 |
| Lysis | Season | Precip | 1.19 | 0.0996 | 0.080 |
| Lysis | Season | R.mass | 1.13 | 0.075 | 0.096 |
| Lysis | Season | Body.molt | 0.553 | 0.411 | 0.149 |
| Agglut | Season | Tdiff | -0.163 | 0.0949 | 0.088 |
| PIT54 | Season | Precip | 0.0321 | 0.0074 | 2.54e-05 |
| PIT54 | Season | R.mass | 0.0127 | 0.00571 | 0.027 |
| PIT54 | Season | Tdiff | 0.0125 | 0.00718 | 0.084 |
| WBC | Year | Tdiff | 0.00686 | 0.00368 | 0.064 |
| WBC | Season | Tdiff | 0.00739 | 0.00371 | 0.048 |
| Lymp | Season | Precip | -0.0206 | 0.0135 | 0.129 |
| Lymp | Season | Tdiff | -0.0181 | 0.0124 | 0.144 |
| Mono | Year | Tmin | -0.00487 | 0.00376 | 0.197 |
| Mono | Season | Tdiff | 0.0114 | 0.00724 | 0.116 |
| Mono | Season | Ff | 0.0294 | 0.0208 | 0.160 |
| Mono | Season | Tmin | 0.00569 | 0.00422 | 0.179 |

Table S7: Lysis: Type II Anova. Horizontal line shows separate models. Model p-value: 1.23e-14 (Year), 9.26e-05 (Season)..

| Time | Covariate | Df | LR Chisq | Pr(>Chisq) |
|---|---|---|---|---|
| Year | Cone.year | 3 | 65.321 | 4.28e-14 |
| Year | Ff | 1 | 3.176 | 0.0747 |
| Season | Tdiff | 1 | 5.400 | 0.0201 |
| Season | Precip | 1 | 3.291 | 0.0697 |
| Season | R.mass | 1 | 2.880 | 0.0897 |
| Season | Body.molt | 1 | 2.083 | 0.149 |
| Season | Season | 3 | 4.858 | 0.183 |
| Season | Ff | 1 | 0.463 | 0.496 |
| Season | Tmin | 1 | 0.081 | 0.776 |

Table S8: Agglut: Type II Anova. Horizontal line shows separate models. Model p-value: 1.84e-11 (Year), 6.21e-04 (Season)..

| Time | Covariate | Df | F value | Pr(>F) |
|---|---|---|---|---|
| Year | Cone.year | 3 | 20.839 | 1.16e-11 |
| Year | Cp.bp | 1 | 0.919 | 0.339 |
| Year | Tdiff | 1 | 0.242 | 0.623 |
| Year | Age | 1 | 0.015 | 0.904 |
| Year | Tmin | 1 | 0.003 | 0.953 |
| Year | Residuals | 183 | | |
| Season | Tdiff | 1 | 2.935 | 0.0884 |
| Season | Tmin | 1 | 0.688 | 0.408 |
| Season | Fat | 1 | 0.500 | 0.481 |
| Season | Season | 3 | 0.094 | 0.964 |
| Season | Residuals | 175 | | |

Table S9: PIT54: Type II Anova. Horizontal line shows separate models. Model p-value: 2.45e-02 (Year), 1.33e-04 (Season)..

| Time | Covariate | Df | F value | Pr(>F) |
|---|---|---|---|---|
| Year | Cone.year | 2 | 5.317 | 0.00587 |
| Year | Body.molt | 1 | 1.272 | 0.261 |
| Year | Sex | 2 | 0.863 | 0.424 |
| Year | Cp.bp | 1 | 0.282 | 0.596 |
| Year | Residuals | 151 | | |
| Season | Precip | 1 | 18.799 | 2.54e-05 |
| Season | R.mass | 1 | 4.980 | 0.027 |
| Season | Tdiff | 1 | 3.013 | 0.0845 |
| Season | Tmin | 1 | 0.586 | 0.445 |
| Season | Season | 3 | 0.518 | 0.67 |
| Season | Fat | 1 | 0.178 | 0.674 |
| Season | Residuals | 162 | | |

Table S10: Lymp: Type II Anova. Horizontal line shows separate models. Model p-value: 1.98e-06 (Year), 1.01e-05 (Season)..

| Time | Covariate | Df | F value | Pr(>F) |
|---|---|---|---|---|
| Year | Cone.year | 3 | 10.093 | 3.75e-06 |
| Year | Precip | 1 | 1.431 | 0.233 |
| Year | Tdiff | 1 | 0.508 | 0.477 |
| Year | Fat | 1 | 0.293 | 0.589 |
| Year | Age | 1 | 0.153 | 0.696 |
| Year | Residuals | 169 | | |
| Season | Season | 3 | 3.537 | 0.0164 |
| Season | Precip | 1 | 2.334 | 0.129 |
| Season | Tdiff | 1 | 2.155 | 0.144 |
| Season | Ff | 1 | 1.622 | 0.205 |
| Season | Sex | 2 | 1.134 | 0.325 |
| Season | Body.molt | 1 | 0.720 | 0.398 |
| Season | Age | 1 | 0.634 | 0.427 |
| Season | Cp.bp | 1 | 0.469 | 0.495 |
| Season | Tmin | 1 | 0.003 | 0.959 |
| Season | Residuals | 144 | | |

Table S11: Mono: Type II Anova. Horizontal line shows separate models. Model p-value: 5.58e-10 (Year), 1.87e-06 (Season)..

| Time | Covariate | Df | F value | Pr(>F) |
|---|---|---|---|---|
| Year | Cone.year | 3 | 19.921 | 4.36e-11 |
| Year | Sex | 2 | 2.100 | 0.126 |
| Year | Tmin | 1 | 1.678 | 0.197 |
| Year | Age | 1 | 1.392 | 0.24 |
| Year | Fat | 1 | 0.746 | 0.389 |
| Year | R.mass | 1 | 0.109 | 0.742 |
| Year | Residuals | 167 | | |
| Season | Season | 3 | 6.146 | 0.000578 |
| Season | Tdiff | 1 | 2.498 | 0.116 |
| Season | Ff | 1 | 1.991 | 0.16 |
| Season | Tmin | 1 | 1.823 | 0.179 |
| Season | R.mass | 1 | 1.109 | 0.294 |
| Season | Sex | 2 | 0.295 | 0.745 |
| Season | Body.molt | 1 | 0.076 | 0.783 |
| Season | Residuals | 146 | | |

Table S12: WBC: Type II Anova. Horizontal line shows separate models. Model p-value: 2.89e-01 (Year), 1.71e-01 (Season)..

| Time | Covariate | Df | F value | Pr(>F) |
|---|---|---|---|---|
| Year | Tdiff | 1 | 3.469 | 0.0643 |
| Year | Cone.year | 3 | 0.676 | 0.568 |
| Year | Tmin | 1 | 0.311 | 0.578 |
| Year | Body.molt | 1 | 0.204 | 0.652 |
| Year | Residuals | 170 | | |
| Season | Tdiff | 1 | 3.963 | 0.0483 |
| Season | Season | 3 | 0.305 | 0.822 |
| Season | Residuals | 156 | | |

### Supplemental Figures

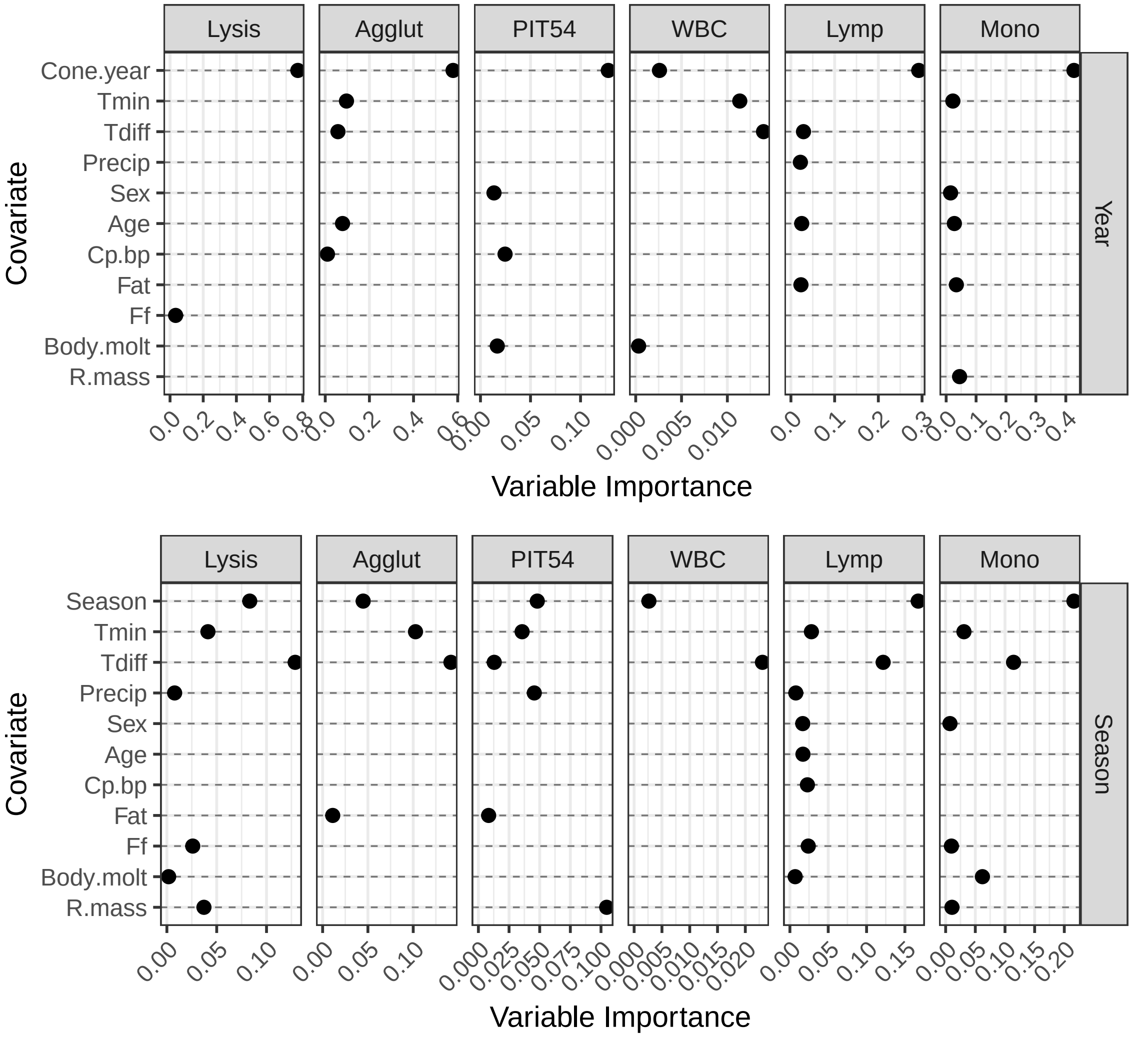


Figure S1: Variable importance for each Random Forest model (RFM, one model per response x time). Covariates with negative variable importance were subsequently removed over 3 serial iterations of model fitting, so that the remaining covariates consistently improve RFM predictive power. To ease comparisons between models, raw importance scores are scaled by model MSE. See Table S4 for model $R^2$s.

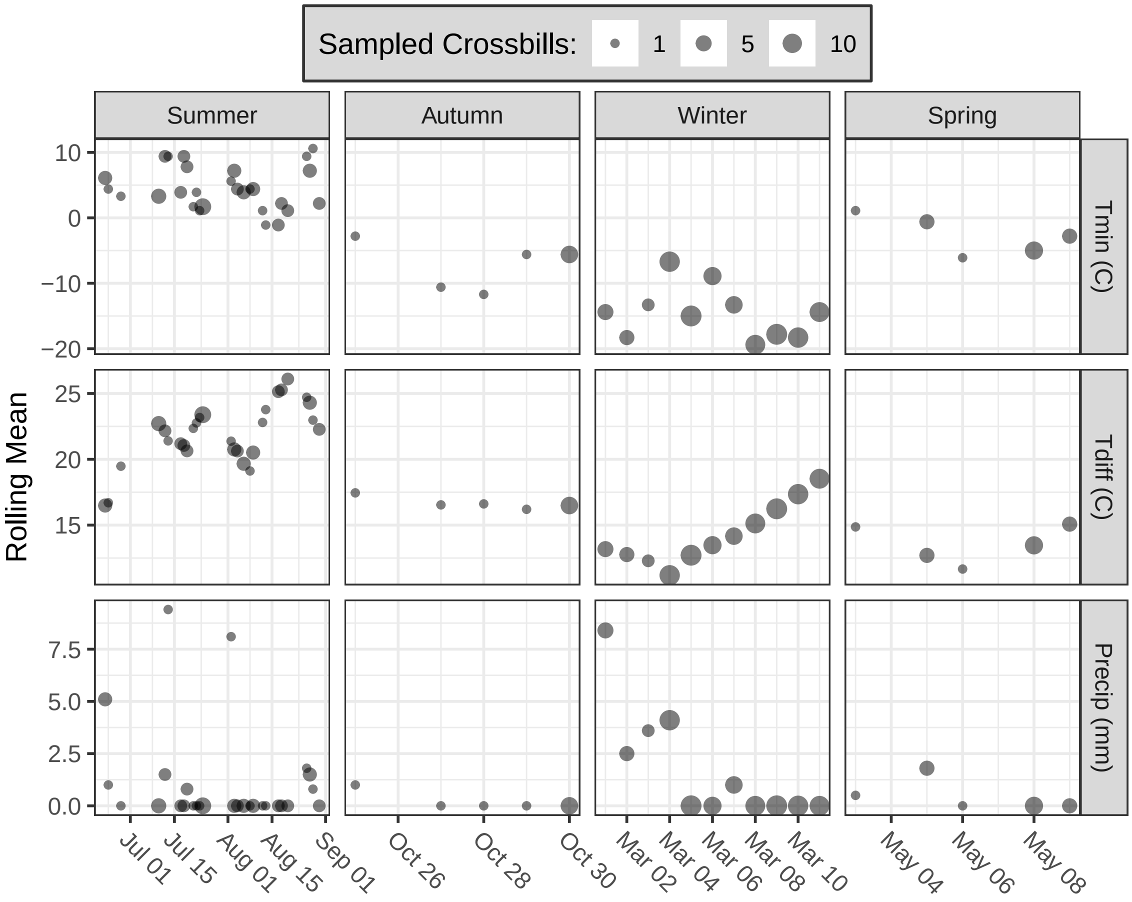


Figure S2: Rolling mean of weather covariates by time, cone year 2011. Point size shows number of captured birds per observation day. See Table S5 for rolling mean details.

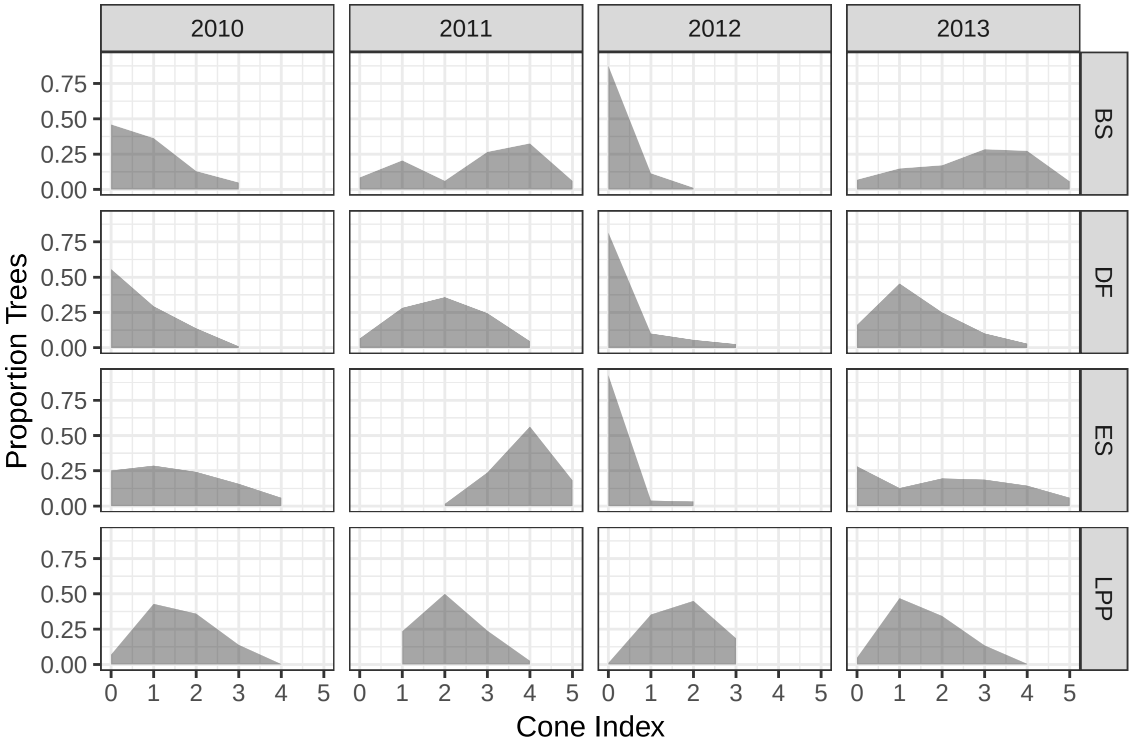


Figure S3: Distribution of cone score by tree species and cone year. Species include blue spruce (BS, *Picea pungens*), Douglas fir (DF, *Pseudotsuga menziesii*), Engelmann spruce (ES, *Picea engelmannii*), and lodge pole pine (LPP, *Pinus contorta*).

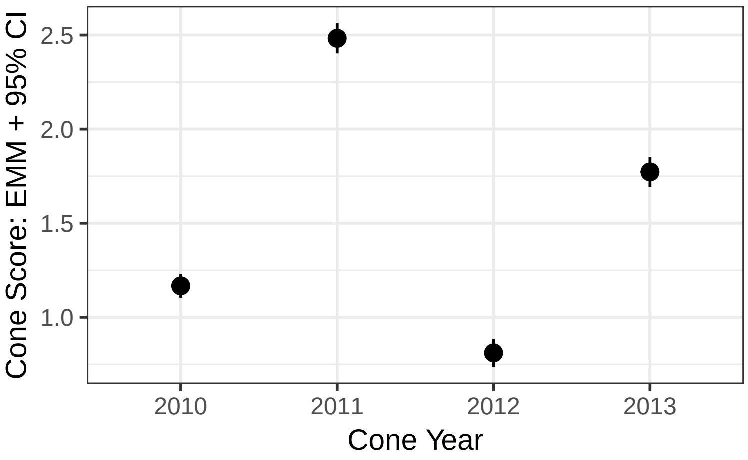


Figure S4: Expected Marginal Means (EMM) of cone score by cone year, showing 95% CI. EMM are computed from an additive linear model of cone score by cone year and tree species (no interactions). All years differ significantly.

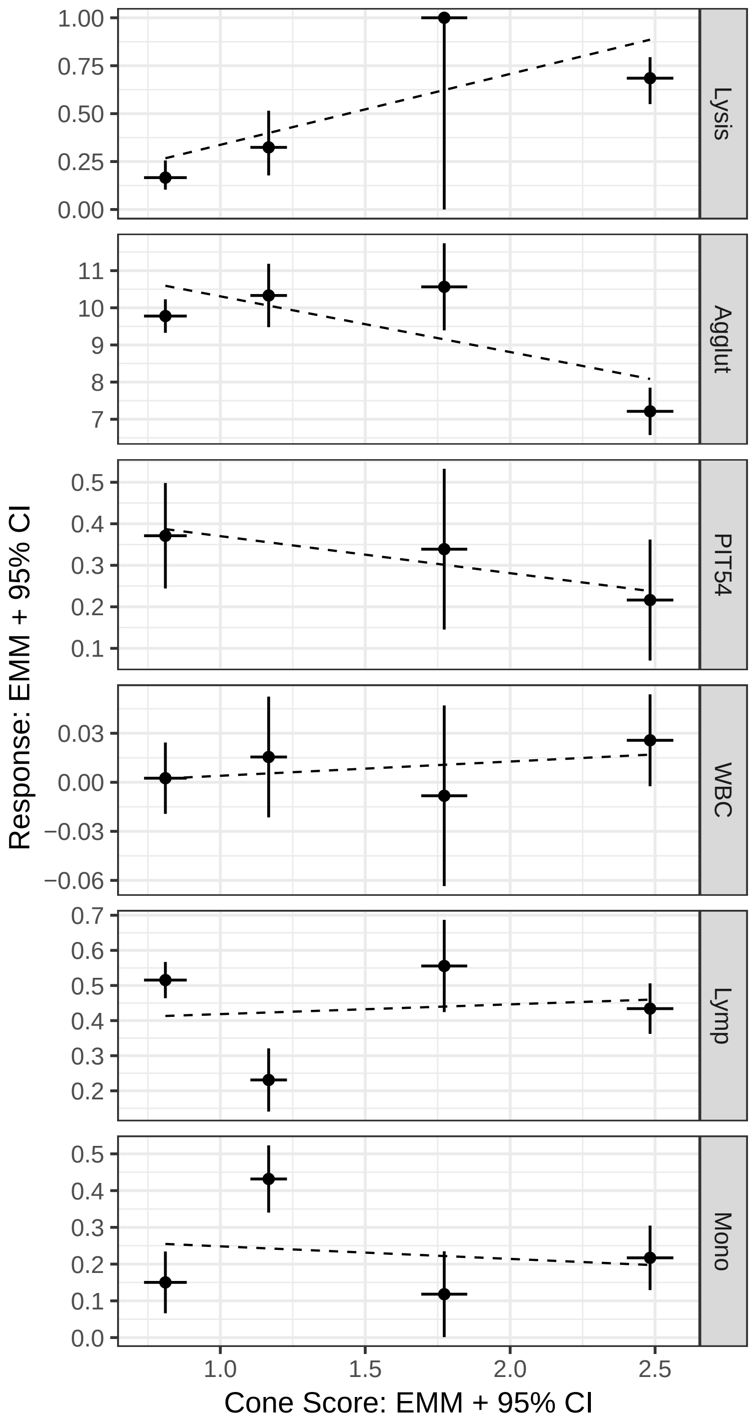


Figure S5: Scatterplot of response EMM versus cone score EMM, for each response. Solid horizontal and vertical lines show respective 95% CI; dashed line shows a best-fit line to respective EMM. See Fig. **??** and Fig. S4 for details.

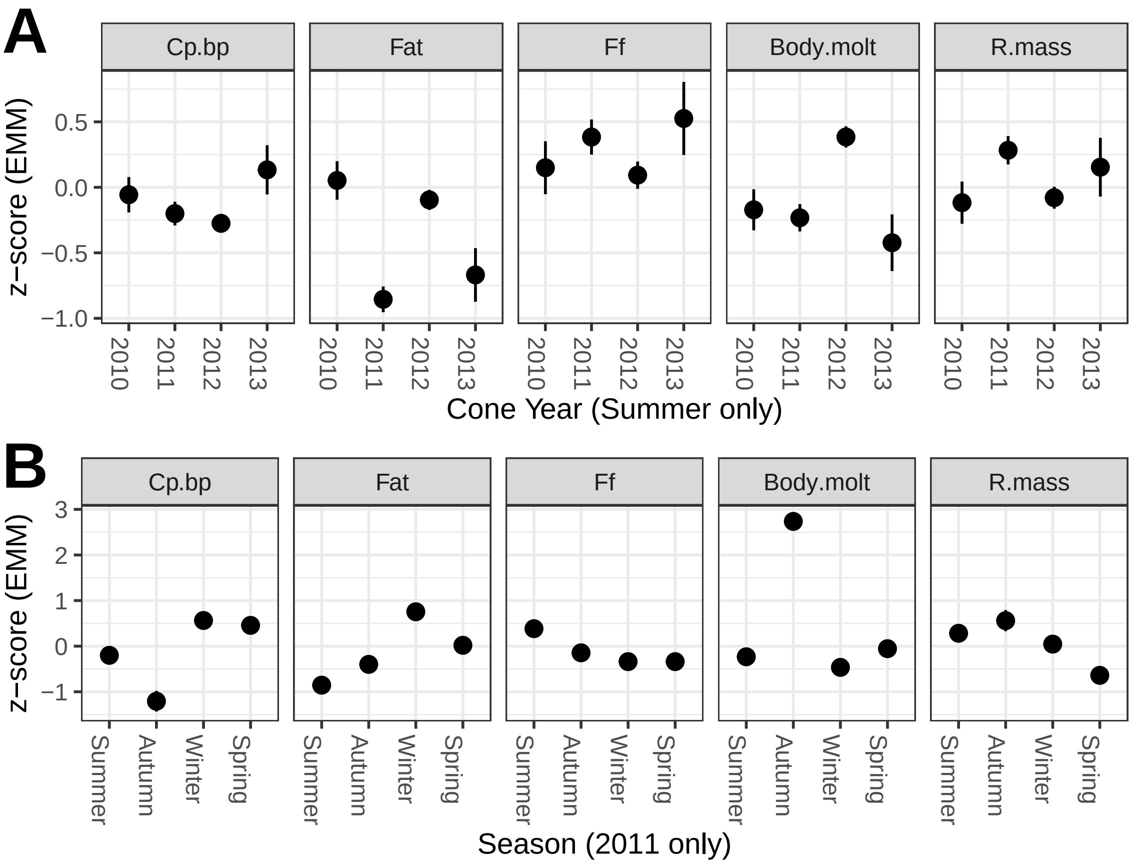


Figure S6: Dependence of continuous physiological covariates on **A**, cone year (Summer observations only) and **B**, season (Cone Year 2011 observations only). Each covariate was z-transformed, and a linear model was constructed using cone year (A) or season (B) as the predictor. The expected marginal mean (EMM) z-score within year (A) and within season (B) are shown, along with 95% CI.

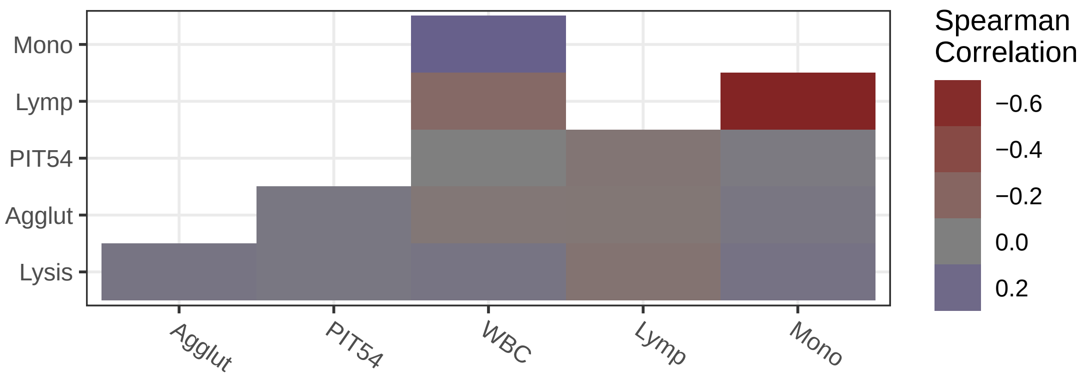


Figure S7: Correlation between measured responses: full period of record.

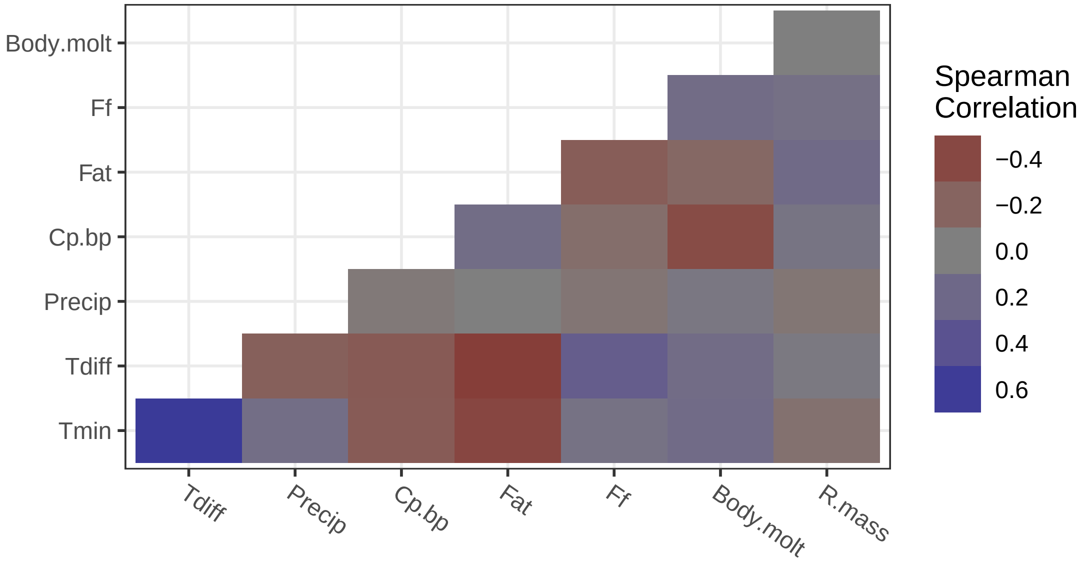


Figure S8: Correlation between covariates: full period of record.

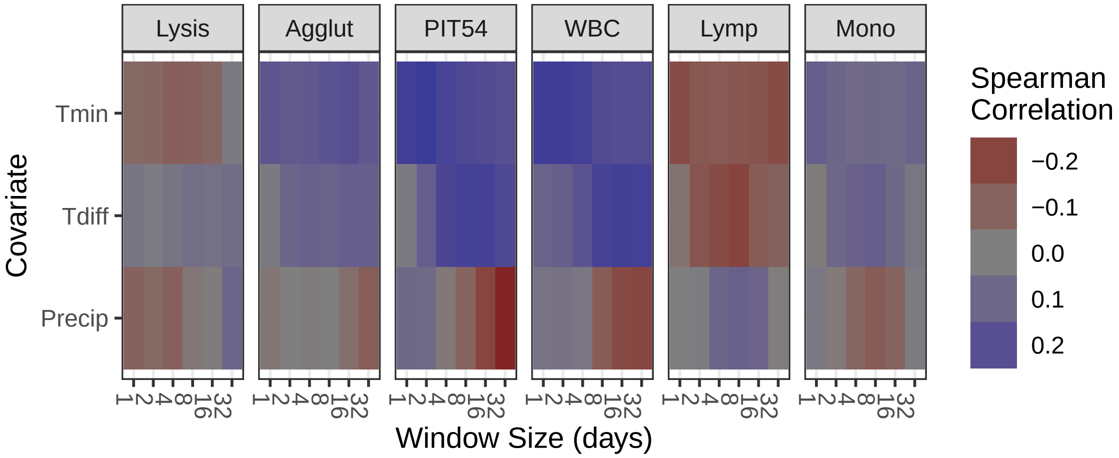


Figure S9: Correlation between response and rolling mean of weather covariates, over a range of window sizes. In general, correlation changes smoothly as window size increases. See also Table S5.
